# Gene-by-environment interactions in agricultural pest management: population effects on diet-*Bt* interactions in a caterpillar

**DOI:** 10.1101/2020.10.30.361170

**Authors:** Carrie Deans, Gregory Sword, Spencer Behmer, Eric Burkness, Marianne Pusztai-Carey, W.D. Hutchison

**Affiliations:** Department of Entomology, University of Minnesota, 219 Hodson Hall, St. Paul, MN 55108; Department of Entomology, Texas A&M University, TAMU 2475, College Station, TX 77843; Department of Biochemistry, Case Western Reserve University, 10900 Euclid Ave., Cleveland, OH 44106

## Abstract

Given that plant nutrient content is both spatially and temporally dynamic (Lenhart et al., 2015; Deans et al., 2016, 2018), insect herbivores are exposed to an incredible amount of nutritional variability. This variability can constrain insects to feeding on sub-optimal resources, but it can also provide an opportunity for insects to regulate their intake of specific nutrients to obtain an optimal balance. Nutrient regulation has implications for pest control strategies in agricultural systems, as the nutritional state of pest species may impact their susceptibility to insecticides. Deans et al. (2017) showed that diet macronutrient balance has significant effects on the susceptibility of *Helicoverpa zea* larvae to Cry1Ac, an endotoxin expressed in transgenic *Bt* crops. This was demonstrated using a highly inbred laboratory strain of *H. zea,* limiting the applicability of these results to field populations that encompass greater genetic diversity. In this study, we assessed the impact of field-relevant macronutrient variability on the efficacy of two *Bt* endotoxins, Cry1Ab and Cry1Ac, using three field populations collected from different geographic regions. This was done to further understand the impact of nutritional variability on *Bt* susceptibility and also to determine the relevance of these effects in the field. While we saw limited differences in Cry susceptibility across populations, dietary effects were highly variable. Across populations there were distinct population-level differences in the interactions between Cry concentration and diet, the type of Cry toxin impacted by diet, and the treatment diet that produced optimal survival and performance. These results show that nutrition can have strong impacts on *Bt* susceptibility but also that these impacts are strongly affected by genetic background in *H. zea*. To accurately assess *Bt* susceptibility in the field, including resistance monitoring, bioassay methods should incorporate the appropriate nutritional parameters and be as localized as possible.

## Introduction

Nutritional variability can have strong impacts on animal fitness. At the organismal level, nutritional constraints on physiological function are well established, but the compounding effects of these constraints on population dynamics, community interactions, and ecosystem function are less well-studied. Consumers are not, however, defenseless when it comes to dealing with variations in resource quality. Selective feeding allows consumers to alter their intake of specific nutrients to respond to the various physiological challenges at hand. In this way, nutritional variability acts as both a challenge and an opportunity. The two nutrients with the greatest impact on animal fitness are dietary protein (p) and carbohydrates (c). The quantitative requirements for these two macronutrients are the greatest of any nutrient, thus they have a high propensity to be limiting. Insect herbivores, particularly generalists, experience immense nutritional variability, as plant macronutrient content is notoriously dynamic. Even within an individual plant, soluble protein and digestible carbohydrate concentrations can differ drastically across spatial gradients, such as tissue types, as well as temporal gradients, like plant ontogeny and phenology (Deans et al., 2016; Lenhart et al., 2015). Insects can detect plant amino acid and sugar concentrations via chemoreception, and they actively regulate their intake of protein and carbohydrates (Behmer, 2009) to reach an optimal ratio, termed an intake target. The ability of an insect to reach this intake target in the field can have strong effects on their survival and performance. Simpson and Raubenheimer (2001) showed that locusts tolerated tannic acid, a grass allelochemical, best when fed on a diet matching their intake target. Additionally, an insect’s optimal p:c ratio may diverge from the intake target under certain circumstances. Behmer et al. (2002) showed that locusts were more willing to ingest tannic acid when it was available in a protein-biased versus a carbohydrate-biased diet. Other kinds of challenges can also be mitigated through nutritional means. Several studies have shown that increases in dietary protein improve immune function in caterpillars challenged with infection and that infected larvae typically regulate for a more protein-biased intake (Povey et al., 2009; Cotter et al., 2010; Shikano and Cory, 2015).

Because nutrition has such strong impacts on animal fitness, it has important implications the management of ecological and agricultural resources. Polyphagy, or the ability to utilize hosts from many different plant families, is a trait commonly found among herbivorous invasive species and insect pests (Cho et al., 2008; Jeschke and Strayer, 2008; Nahrung and Swain, 2015; Pearce et al., 2017). Adaptations associated with polyphagy include the ability to tolerate a diversity of plant secondary compounds in addition to nutritional variability; however, these two traits are likely intimately connected, given that detoxification is fundamentally limited by nutritional constraints (Lindroth et al., 1990, 1991; Patil et al., 1990; Berenbaum and Zangerl, 1994; Jensen et al., 2016). Nutritional plasticity allows invasive and pest species to utilize wider host ranges and invade new habitats. Additionally, their physiological flexibility may also limit the effectiveness different management tactics. In this study, we explore the relationship between nutritional plasticity and toxicity in an agricultural system to show the importance of understanding gene-by-environment interactions and to highlight the value of this understanding for resource management.

The adoption of transgenic *Bt* crops has been widespread since its introduction in 1996. Currently, over 75% of U.S. corn and cotton acreage are comprised of varieties containing at least one *Bt* gene (www.ers.usda.gov/publications/erreconomic-researchreport/err162.aspx). These varieties contain a gene or genes that encode endotoxins indigenous to the soil bacteria *Bacillus thuringiensis*. These *Bt* toxins, which are referred to as Cry endotoxins, are proteins that have insecticidal properties in the gut environment of specific lepidopteran and coleopteran insect species, particularly heliothine moths, European corn borer, and corn rootworm. *Bt* spore solutions are also used extensively as an infectious topical agent in organic farming. In 2017, Deans et al. showed that field-relevant variation in dietary macronutrient profiles could significantly alter the susceptibility of *Helicoverpa zea* larvae to the *Bt* toxin Cry1Ac. Results showed that larvae reared on an artificial diet approximating the intake target exhibited the best survival when challenged with Cry1Ac and that those reared on either a carbohydrate- or protein-biased diet had significantly lower survival. Also, when susceptibility was assessed using a dose-response bioassay, the standard method used in resistance monitoring, the median lethal concentration for Cry1Ac was increased by over 100-fold by simply changing the artificial diet from a carbohydrate-biased commercial diet to one with a p:c ratio matching the intake target. This study showed the potential for field-relevant nutritional variation to impact *Bt* efficacy; however, these studies were performed using a highly-inbred laboratory strain of *H. zea*, making the applicability of these results to natural, more genetically diverse *H. zea* populations more limited. Opert et al. (2015) found that diet p:c ratio had no effect on *Bt* susceptibility for a resistant strain of *H. zea* but did have an effect on the susceptible parental strain. Additionally, Shikano and Corey (2014) found differences in the effect of diet p:c ratio on *Bt* toxicity in a susceptible and resistant strain of *Trichoplusia ni*. Although both studies utilized spore/pro-toxin as the source of *Bt*, rather than pure trypsin-activated toxin indicative to transgenic *Bt* plants, their results do suggest that genetic differences can affect the impact of nutrition on *Bt* susceptibility.

In order to determine the importance of nutritionally-mediated impacts on *Bt* susceptibility in the field, we assessed the effect of field-relevant macronutrient variation on susceptibility to two different *Bt* endotoxins using three geographically-separate populations of *H. zea*. Susceptibility to Cry1Ac and Cry1Ab, which have been implemented in *Bt* cotton and corn since the 1990s, was initially assessed for all field populations using dose-response bioassays. These baseline data were then used to calculate Cry concentrations at standardized lethal doses (controls, LC25, and LC75) for each population. Survival and performance were measured for larvae reared on one of three different diets simulating the nutrient profile of sweet corn (0.4:1), developing cotton seed (1.6:1), and cotton leaves (2.4:1) (Figure 2) with Cry incorporated. We expected that each population would show differences in initial susceptibility due to genetic distinctions between field populations and that any differential responses to diet across populations would indicate gene-by-environment interactions related to nutritionally-mediated susceptibility. Understanding the prevalence of nutritionally-mediated impacts on *Bt* susceptibility will help to identify factors that contribute to regional variability in *Bt* efficacy, and will inform aspects of insect biology that may be exploited to improve these biotechnologies. Documenting these nutritional effects is also important for optimizing resistance monitoring by make sure the most field-relevant diets are being used in diet-incorporated bioassays.

## Methods

### Insect Colonies

Three different colonies of *H. zea* were created and maintained to supply eggs for subsequent experiments. Figure 1 shows the geographic locations where each colony originated: Minnesota (UMORE Agricultural Research Station, Rosemount, MN), Louisiana (LSU Kerns Lab, Winnsboro, LA), and Missouri (USDA Little Lab, Stoneville, MS). The LA and MS colonies were started from eggs provided by collaborator’s pre-existing colonies. The MN population was started from larvae collected from late-season sweet corn in 2016. All colonies were housed in a single walk-in growth chamber (27°C, 14:10 L:D). All colonies were maintained on a soy-based artificial diet (Southland Products Inc.: Lake Village, AR) during the larval stage and on 10% sucrose solutions as adults. For each population, all adults were allowed to mate and lay eggs in large mesh cages. Between 200-400 neonates were randomly collected from each population cage and placed on diet in trays with individual cells to supplement each colony.

**Figure 1.**
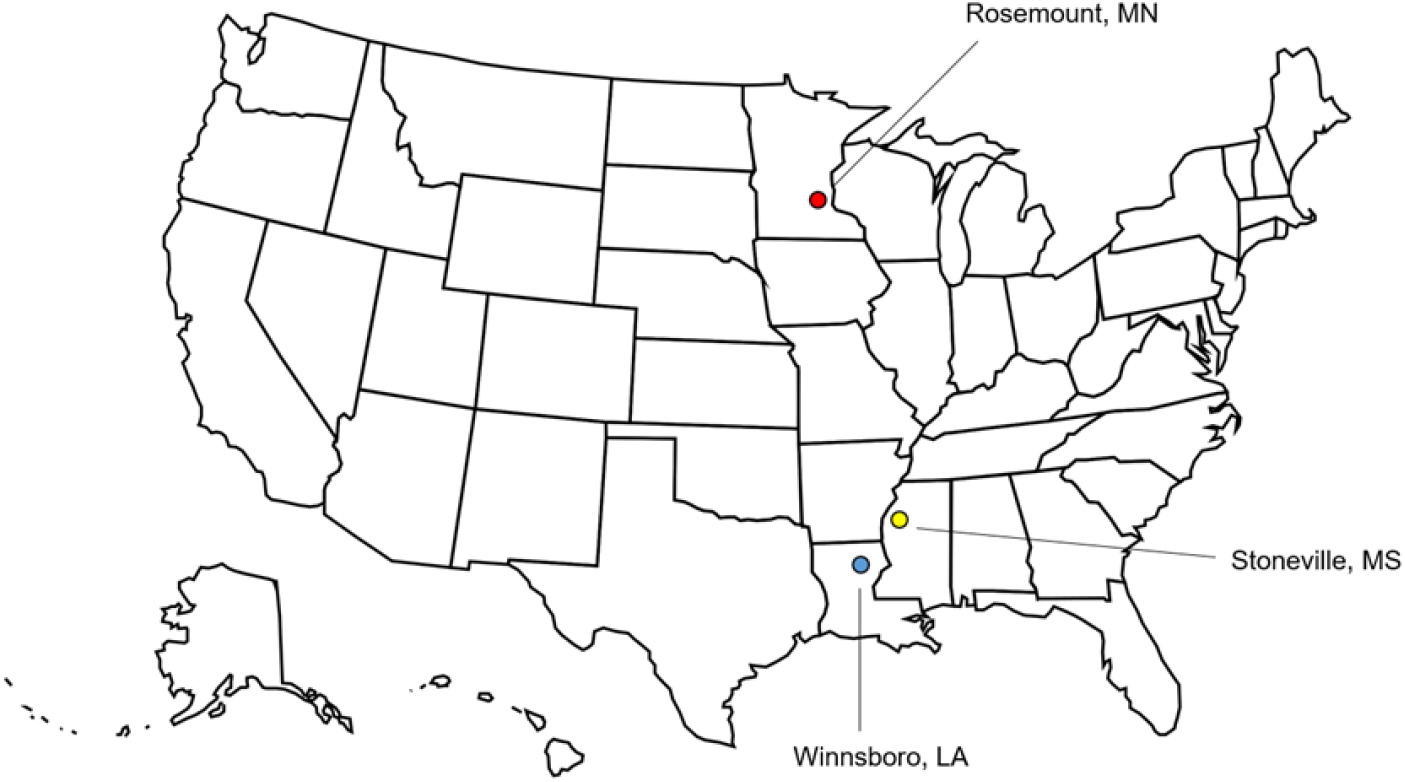
Map of the locations where each of the three field populations came from.

**Figure 2.**
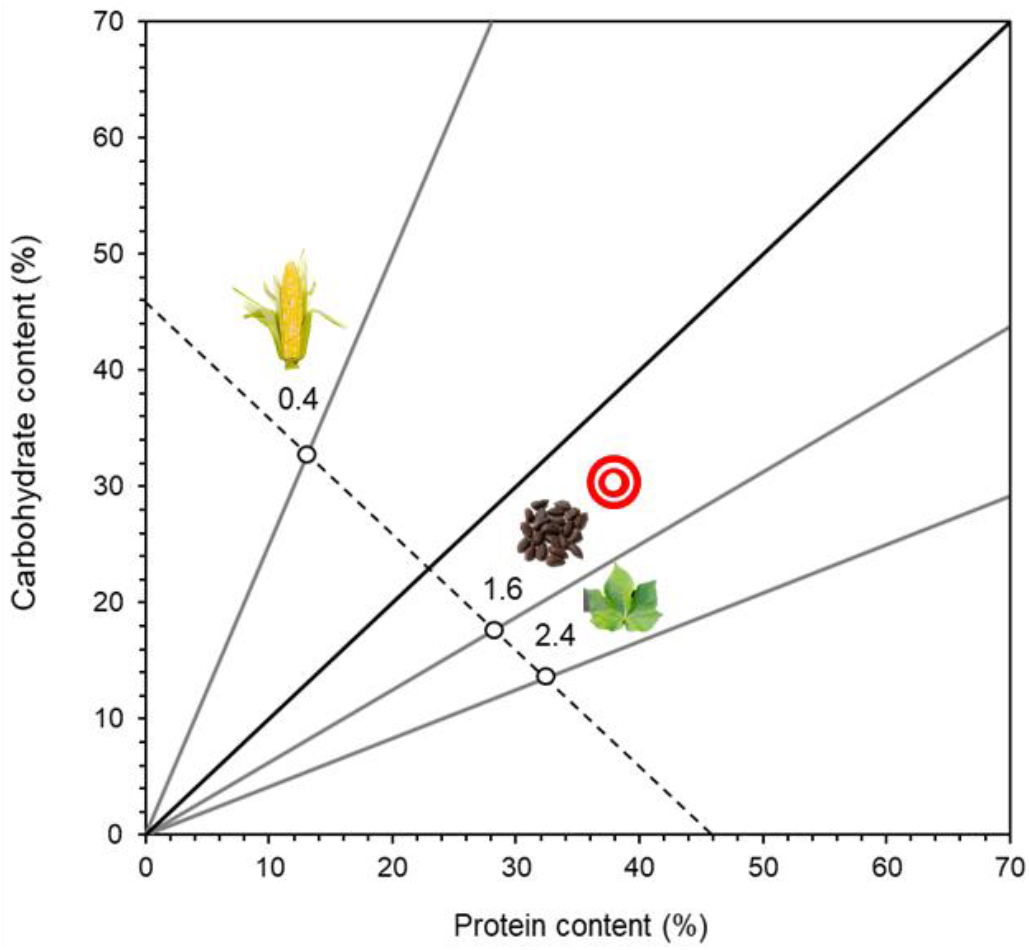
A bi-coordinate plot showing the p:c ratio and p+c concentration for the three diets tested. The black line represents a 1:1 ratio. Gray lines represent p:c ratios typifying sweet corn kernels (0.4), developing cotton seed (1.6), and cotton leaves (2.4). The target also shows the diet matching the reported intake target for *H. zea*.

### Experimental Artificial Diets

The artificial diet described in Jing et al. (2013) was altered to create the experimental diets for this study. Based largely on the diet of Ritter and Nes (1981), the key ingredients in the Jing et al. (2013) diet are vitamin-free casein, sucrose, cellulose, Wesson’s salt mix, Torula yeast, lipids (cholesterol, linoleic and linolenic acid) and vitamins. This diet has been shown to produce high survival and to support successful development in *H. zea* (Jing et al., 2013; Deans et al., 2015; Deans et al., 2017). A small amount of the colony maintenance diet (Southland diet) (an amount commensurate to 20% of total dry mass) was added to the recipe, and reductions in agar, methyl paraben, sorbic acid, and chloro-tetracycline were made to account for the addition of these ingredients from the Southland diet, thereby maintaining the percentage of these ingredients at the same levels found in the original diet. Only three ingredients were altered to achieve the macronutrient profiles described below: cellulose, casein (primary protein source), and sucrose (primary carbohydrate source). All other ingredients were consistent across experimental diets.

Plant macronutrient data for cotton (Deans et al., 2016) and sweet corn (Deans et al., 2018) were used to select field-relevant macronutrient profiles to model the experimental diet treatments after. *H. zea* is a notorious pest of cotton and sweet corn and additionally, transgenic *Bt* cotton and sweet corn varieties are routinely planted throughout the U.S. to control *H. zea* populations. Figure 2 shows the three diets that were created to represent a range of p:c ratios; 0.4 (13p:33c), 1.6 (p26:c20), and 2.4 (p32.5:c13.5). All diets contained a total macronutrient content of 46%, a value that is common among across cotton and sweet corn tissues (Deans et al., 2016, 2018). The 0.4 diet, which is carbohydrate-biased, represents the profile of sweet corn kernels (Deans et al., 2018), while the 1.6 diet matches that of the self-selected intake target for *H. zea* (Deans et al., 2015) as well as the ratio of developing cotton seed. Lastly, the protein-biased 2.4 diet is indicative of cotton leaves (Deans et al., 2016).

### Cry Bioassays

Once each colony was established, dose-response bioassays for Cry1Ac and Cry1Ab were performed for each population to determine baseline susceptibility. Cry proteins were incorporated into the experimental artificial diet (see below) matching the self-selected intake target (p:c of 1.6:) for *H. zea* (Deans et al., 2015) at seven concentrations: 0, 0.01, 0.1, 1, 10, 100, 1000 ppm. Trypsin-activated HPLC-purified Cry1Ac and Cry1Ab was procured from the Pusztai-Carey Lab at Case Western Reserve University (Cleveland, OH) and stored at −80°C. Stock solutions were made for each Cry concentration tested such that the amount of solution, and thus the overall volume of liquid, added to the diet was standardized for each treatment. Stock solutions were stored at −20°C and thawed for diet preparation. The required amount of stock solution needed to meet the desired concentration (ppm) was calculated based on the total wet mass of diet being deployed in each treatment (ug of Cry per g of diet). Distilled water was added to the 0 ppm treatments in the same volume added to the other treatments. The diet was thoroughly mixed with the Cry solution before being portioned into single-cell rearing trays. Neonates were then individually placed on diet in each treatment (N=20-32) and mortality was measured after 10 days. A probit analysis was used to create dose-response curves for each Cry protein for each population. In particular, the LC_25_, LC_50_, and LC_75_ values were calculated for each population and Cry protein to inform the treatment doses selected for the full diet experiment (see below).

### Cry-Diet Experimental Protocol

For each population, three Cry1Ac and Cry1Ab concentrations (no Cry, a LC_25_ dose, and a LC_75_ dose) were tested on each of the three experimental diets (0.4, 1.6, and 2.4), resulting in a total of 12 different treatments tested per Cry protein per population. The specific Cry doses were determined by the results of the initial Cry bioassays. As a result, the exact concentrations of Cry1Ac and Cry1Ab tested did vary across populations, but because each dose corresponded to the same lethal concentration (LC_25_, LC_75_), the physiological impact of Cry was standardized across populations for each treatment. For each population, stock solutions of Cry1Ac and Cry1Ab were made, stored at −20°C, and thawed for diet preparation. As with the *Bt* bioassays, the total volume of solution added to diet in each Cry treatment was standardized. Diets and solutions were mixed thoroughly, diet was placed into single-cell trays with perforated sticker lids, and one neonate per cell was deployed (N=18-32). Individuals were followed through eclosion. Survival, time to pupation (developmental time), pupal mass, and sex were recorded.

### Data Analysis

Each population was tested at a different time, precluding direct statistical comparisons between populations; however, Cry1Ac and Cry1Ab trials within each population were run concurrently. Because of this, statistics for each population were performed separately. A Kaplan-Meier analysis was used to determine significant differences in mortality across Cry treatments and diets and also differences in developmental time between Cry concentrations and diet treatments. An ANOVA was used to test for differences in pupal mass across treatments. All statistics were done using IBM SPSS Statistics v.22.

## Results

### Cry Bioassays

Table 1 shows the LC_25_, LC_50_, and LC_75_ values for Cry1Ac and Cry1Ab for each population, as well as the %95 confidence intervals. The MN population exhibited the lowest values across all lethal concentrations as well as overlapping CIs for both Cry1Ac and Cry1Ab, indicating similar susceptibilities for both endotoxins. The MS population showed slightly higher lethal concentrations that were between 1.2-2.2 times higher than that of the MN Cry1Ac values and 2.6-8.2 times higher than the MN Cry1Ab values. The MS population also showed a higher tolerance to Cry1Ab compared to Cry1Ac, as the values for the LC_50_ and LC_75_ values were higher with non-overlapping confidence intervals. The LA population showed the highest lethal concentration values of all the populations tested, surpassing the highest MS Cry1Ac lethal dose by 14.4 times and the highest Cry1Ab lethal dose by as much as 4.2 times. The susceptibility to Cry1Ac and Cry1Ab was similar for the LA population, as there was considerable overlap across confidence intervals for all lethal concentrations.

**Table 1.** Dose-response results for Cry1Ab and Cry1Ac for each of the three field-collected populations.

| Population | Cry | LC<sub>25</sub> | LC<sub>50</sub> | LC<sub>75</sub> |
| --- | --- | --- | --- | --- |
| MN | 1Ac | 0.192 (0.05-0.70) | 1.62 (0.45-5.91) | 13.65 (3.75-49.78) |
|  | 1Ab | 0.078 (0.02-0.34) | 1.33 (0.30-5.79) | 22.75 (5.24-98.76) |
| MS | 1Ac | 0.423 (0.13-1.33) | 2.69 (0.86-8.49) | 17.23 (5.48-54.20) |
|  | 1Ab | 0.644 (0.20-2.08) | 6.22 (19.24-20.11) | 60.11 (18.59-194.37) |
| LA | 1Ac | 0.771 (0.17-3.42) | 13.81 (3.06-61.43) | 247.49 (51.08-1026.70) |
|  | 1Ab | 0.254 (0.07-1.16) | 8.02 (2.21-35.95) | 253.17 (61.11-995.75) |

### Cry-Diet Experiments

#### MN Population

Cry concentration had a significant effect on larval survival for both Cry1Ac and Cry1Ab. Overall, as Cry concentration increased, survival decreased (Figure 3a,d,g; Figure 4a,d,g). The Kaplan-Meier analysis showed that for both Cry endotoxins there were no significant differences in survival between the controls and the LC_25_ treatment; however, there were significant differences between the controls and LC_75_ treatment and the LC_25_ and LC_75_ treatments (Table 2). Survival under control conditions was high, with an average of survival 87.5% across all diets. Based on the results of the initial dose-response bioassays done on the 1.6 diet, survival was much lower than expected for the Cry1Ac LC_25_ treatment, at 41.7%. The LC_75_ treatments also showed higher mortality than expected, as no larvae survived to pupation. For Cry1Ab, survival was only slightly lower than expected in the LC_25_ treatment, with survival at 62.5%; however, LC_75_ treatment survival was again higher than expects at 0%.

**Table 2.**
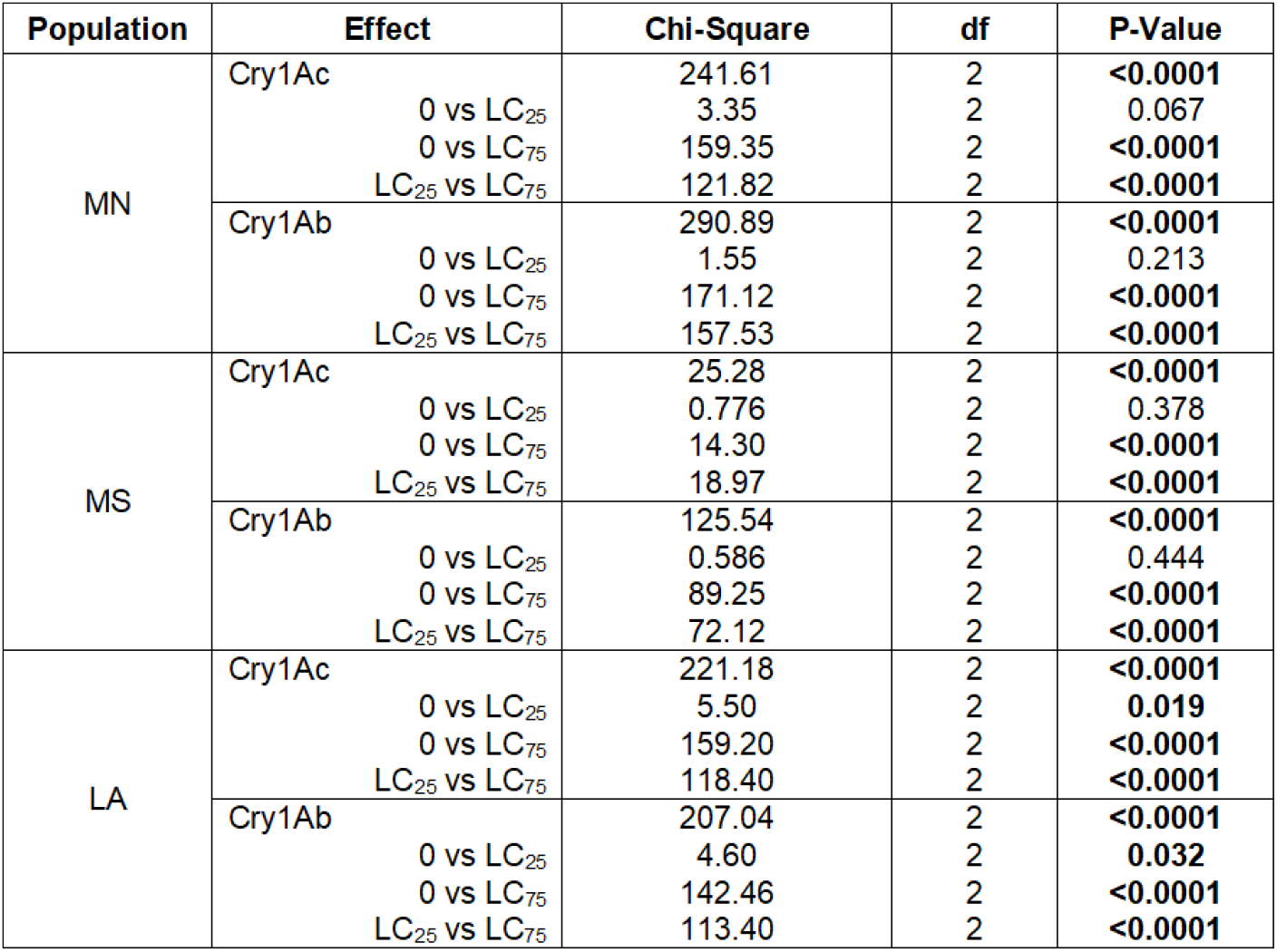
Kaplan-Meier (log rank test) results for the effect of Cry1Ac and Cry1Ab on survival and pairwise comparisons for each population, and Bolded values indicate significance at P ≤ 0.05.

**Figure 3.**
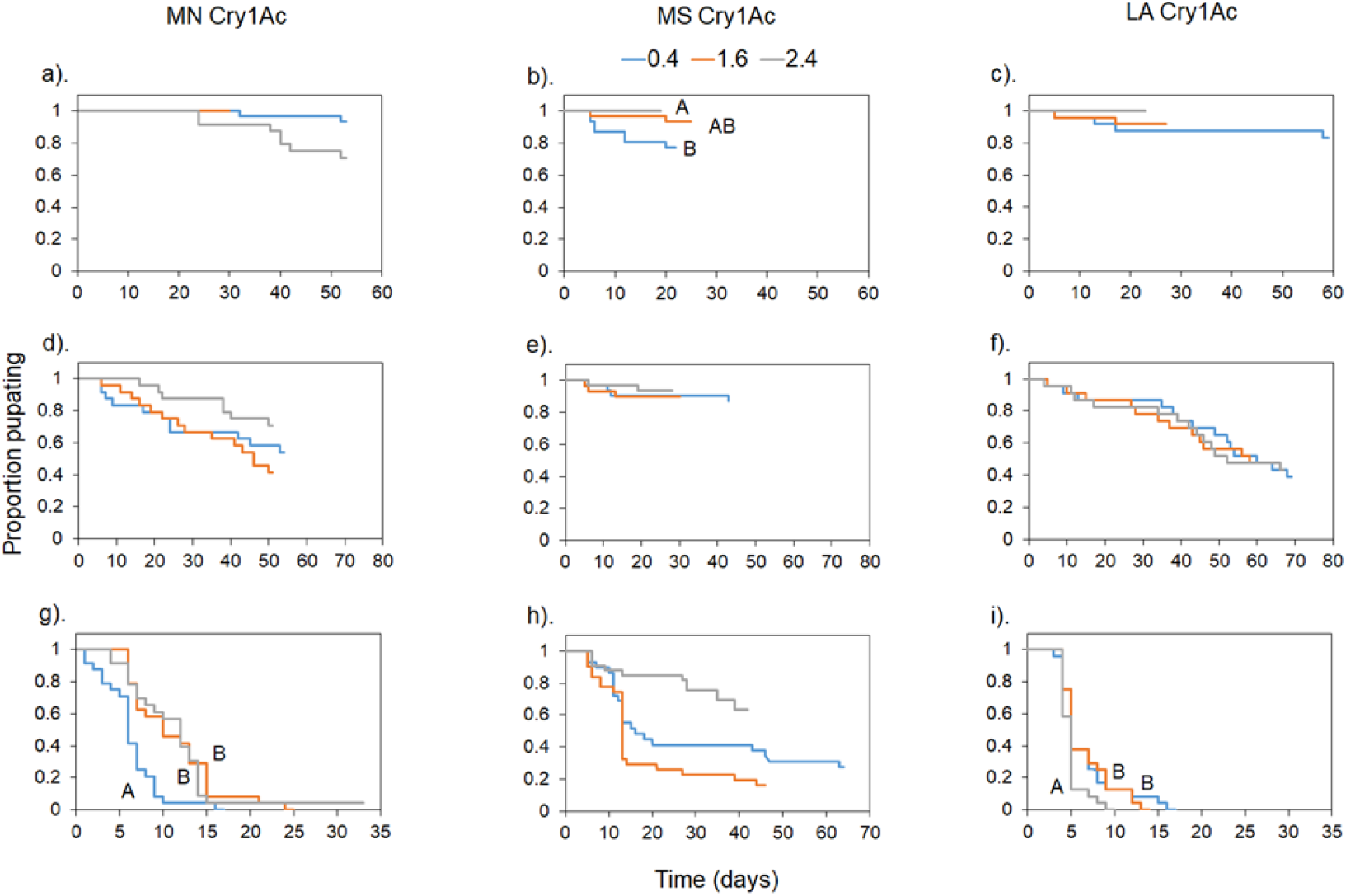
Survival curves for each population (N=18-32) when exposed to no Cry1Ac (a-c), an LC25 dose of Cry1Ac (d-f), and an LC75 dose of Cry1Ac (g-i).

**Figure 4.**
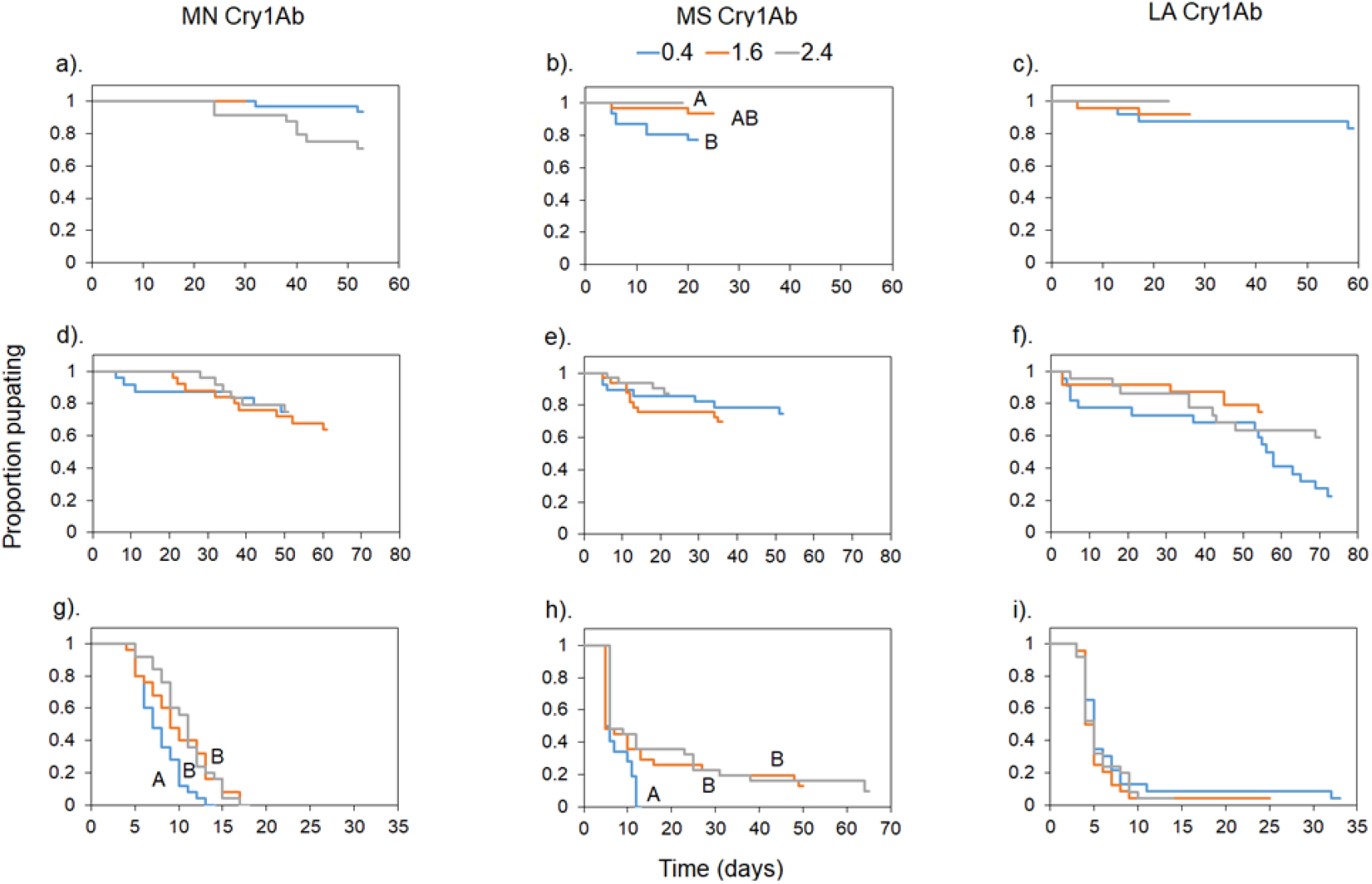
Survival curves for each population (N=18-32) when exposed to no Cry1Ab (a-c), an LC25 dose of Cry1Ab (d-f), and an LC75 dose of Cry1Ab (g-i).

Diet effects were evident and similar for both Cry1Ac and Cry1Ab. There were no significant diet effects on survival for the control and LC_25_ treatments, but there were significant diet effects in the LC_75_ treatments for Cry1Ac and Cry1Ab (Table 3). Figure 3g shows that while all larvae died in the 0.4 and 1.6 treatments, death was quicker for larvae fed the carbohydrate-biased 0.4 diet and mortality curves were similar for the 1.6 and 2.4 diet treatments. Mortality reached 100% for all diets in the Cry1Ab treatments, however, the same statistical pattern was apparent, where larvae on the 0.4 diet died faster than those on either the 1.6 and 2.4 diet (Figure 4g).

**Table 3.** Kaplan-Meier (log rank test) results for the effect of diet on the survival curves for each population within each Cry treatment. Bolded values indicate significance at P ≤ 0.05.

| Population | Effect |  | Chi-Square | df | P-Value |
| --- | --- | --- | --- | --- | --- |
| MN | Cry1Ac- diet | controls | 2.14 | 2 | 0.343 |
|  |  | LC<sub>25</sub> | 2.40 | 2 | 0.310 |
|  |  | LC<sub>75</sub> | 18.79 | 2 | <0.0001 |
|  | Cry1Ab- diet | controls | 2.14 | 2 | 0.343 |
|  |  | LC<sub>25</sub> | 0.174 | 2 | 0.917 |
|  |  | LC<sub>75</sub> | 12.44 | 2 | 0.002 |
| MS | Cry1Ac- diet | controls | 9.44 | 2 | 0.009 |
|  |  | LC<sub>25</sub> | 0.182 | 2 | 0.913 |
|  |  | LC<sub>75</sub> | 3.56 | 2 | 0.168 |
|  | Cry1Ab- diet | controls | 9.44 | 2 | 0.009 |
|  |  | LC<sub>25</sub> | 2.38 | 2 | 0.305 |
|  |  | LC<sub>75</sub> | 9.88 | 2 | 0.007 |
| LA | Cry1Ac- diet | controls | 2.86 | 2 | 0.239 |
|  |  | LC<sub>25</sub> | 1.12 | 2 | 0.572 |
|  |  | LC<sub>75</sub> | 6.05 | 2 | 0.049 |
|  | Cry1Ab- diet | controls | 2.04 | 2 | 0.360 |
|  |  | LC<sub>25</sub> | 0.543 | 2 | 0.762 |
|  |  | LC<sub>75</sub> | 0.568 | 2 | 0.753 |

Cry concentration had significant effects on larval development time for both Cry1Ac and Cry1Ab (Table 4). Figure 5 shows that developmental time was significantly longer in the LC_25_ treatments compared to the controls. Diet had no significant effect on developmental time for the MN population (Table 5). There was a significant effect of Cry concentration on pupal mass for both Cry proteins (Table 6), with pupae weighing significantly less in the LC_25_ treatment than the controls (Figure 6). There was no diet effect on pupal mass (Table 6).

**Table 4.**
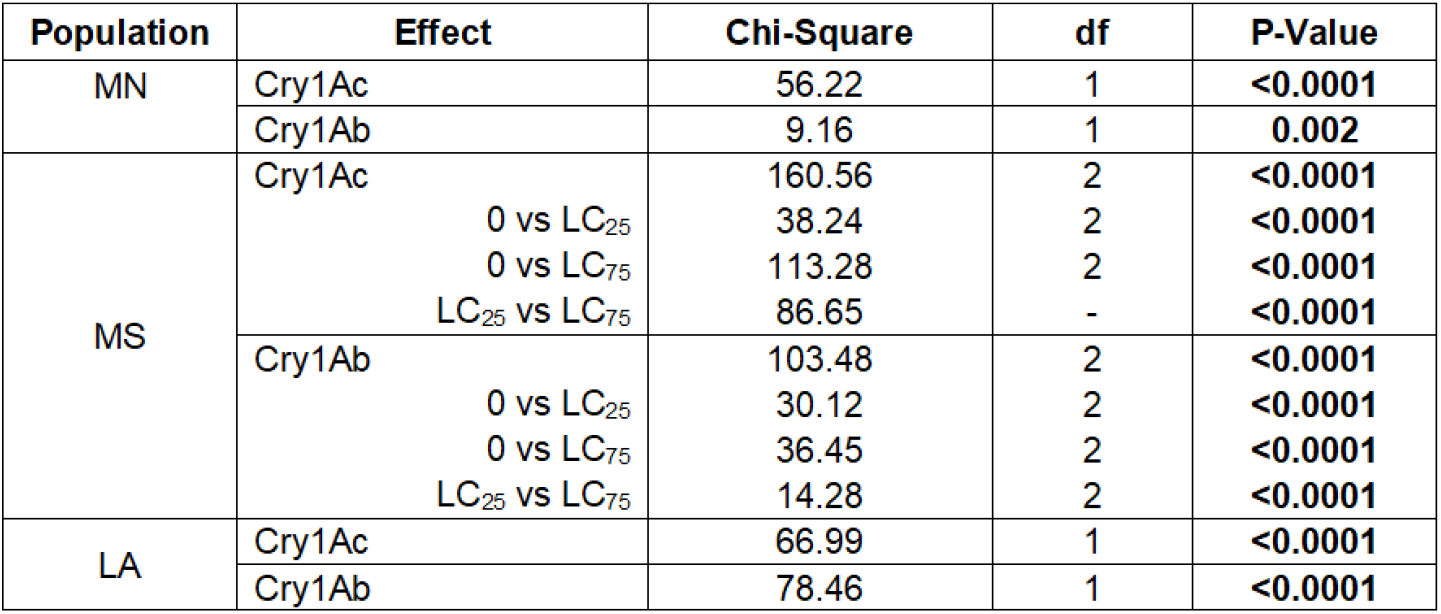
Kaplan-Meier (log rank test) results for the effect of Cry on developmental time for each population. Bolded values indicate significance at P≤ 0.05. The MN and LA populations only had pupation events in the controls and LC_25_ treatments, hence post-hoc results are not listed.

| Population | Effect | Chi-Square | df | P-Value |
| --- | --- | --- | --- | --- |
| MN | Cry1Ac | 56.22 | 1 | **<0.0001** |
|  | Cry1Ab | 9.16 | 1 | **0.002** |
| MS | Cry1Ac | 160.56 | 2 | **<0.0001** |
|  | 0 vs LC<sub>25</sub> | 38.24 | 2 | **<0.0001** |
|  | 0 vs LC<sub>75</sub> | 113.28 | 2 | **<0.0001** |
|  | LC<sub>25</sub> vs LC<sub>75</sub> | 86.65 | - | **<0.0001** |
|  | Cry1Ab | 103.48 | 2 | **<0.0001** |
|  | 0 vs LC<sub>25</sub> | 30.12 | 2 | **<0.0001** |
|  | 0 vs LC<sub>75</sub> | 36.45 | 2 | **<0.0001** |
|  | LC<sub>25</sub> vs LC<sub>75</sub> | 14.28 | 2 | **<0.0001** |
| LA | Cry1Ac | 66.99 | 1 | **<0.0001** |
|  | Cry1Ab | 78.46 | 1 | **<0.0001** |

**Table 5.** Kaplan-Meier (log rank test) results for the effect of diet on developmental time for each population. Bolded values indicate significance at P≤ 0.05. The MN and LA populations only had pupation events in the controls and LC_25_ treatments, hence analyses for the LC_75_ treatments were not performed.

| Population | Effect | Chi-Square | df | P-Value |
| --- | --- | --- | --- | --- |
| MN | Cry1Ac controls | 1.54 | 2 | 0.464 |
|  | LC<sub>25</sub> | 1.46 | 2 | 0.482 |
|  | LC<sub>75</sub> | - | - | - |
|  | Cry1Ab controls | 1.54 | 2 | 0.464 |
|  | LC<sub>25</sub> | 2.95 | 2 | 0.228 |
|  | LC<sub>75</sub> | - | - | - |
| MS | Cry1Ac controls | 30.12 | 2 | **<0.0001** |
|  | LC<sub>25</sub> | 14.94 | 2 | **0.001** |
|  | LC<sub>75</sub> | 4.18 | 2 | 0.124 |
|  | Cry1Ab controls | 30.21 | 2 | **<0.0001** |
|  | LC<sub>25</sub> | 51.39 | 2 | **<0.0001** |
|  | LC<sub>75</sub> | 4.66 | 1 | **0.031** |
| LA | Cry1Ac controls | 7.03 | 2 | **0.030** |
|  | LC<sub>25</sub> | 0.503 | 2 | 0.777 |
|  | LC<sub>75</sub> | - | - | - |
|  | Cry1Ab controls | 7.03 | 2 | **0.030** |
|  | LC<sub>25</sub> | 1.38 | 2 | 0.502 |
|  | LC<sub>75</sub> | - | - | - |

**Table 6.** Two-way ANOVA results for the effect of Cry and diet on pupal mass for that the differences each population. Bolded values indicate significance at P ≤ 0.05.

| Population | Effect | F Stat | df | P-Value |
| --- | --- | --- | --- | --- |
| MN | Cry1Ac | 42.84 | 2 | **<0.0001** |
|  | Diet | 0.850 | 2 | 0.431 |
|  | Cry * Diet | 0.558 | 2 | 0.574 |
|  | Cry1Ab | 28.15 | 1 | **<0.0001** |
|  | Diet | 0.485 | 2 | 0.617 |
|  | Cry * Diet | 0.397 | 2 | 0.674 |
| MS | Cry1Ac | 26.43 | 2 | **<0.0001** |
|  | Diet | 2.63 | 2 | 0.075 |
|  | Cry * Diet | 0.68 | 4 | 0.606 |
|  | Cry1Ab | 19.38 | 2 | **<0.0001** |
|  | Diet | 0.223 | 2 | 0.801 |
|  | Cry * Diet | 0.912 | 4 | 0.437 |
| LA | Cry1Ac | 97.64 | 1 | **<0.0001** |
|  | Diet | 0.889 | 2 | 0.415 |
|  | Cry * Diet | 1.76 | 2 | 0.178 |
|  | Cry1Ab | 24.65 | 2 | **<0.0001** |
|  | Diet | 2.55 | 2 | 0.083 |
|  | Cry * Diet | 487.57 | 2 | 0.416 |

**Figure 5.**
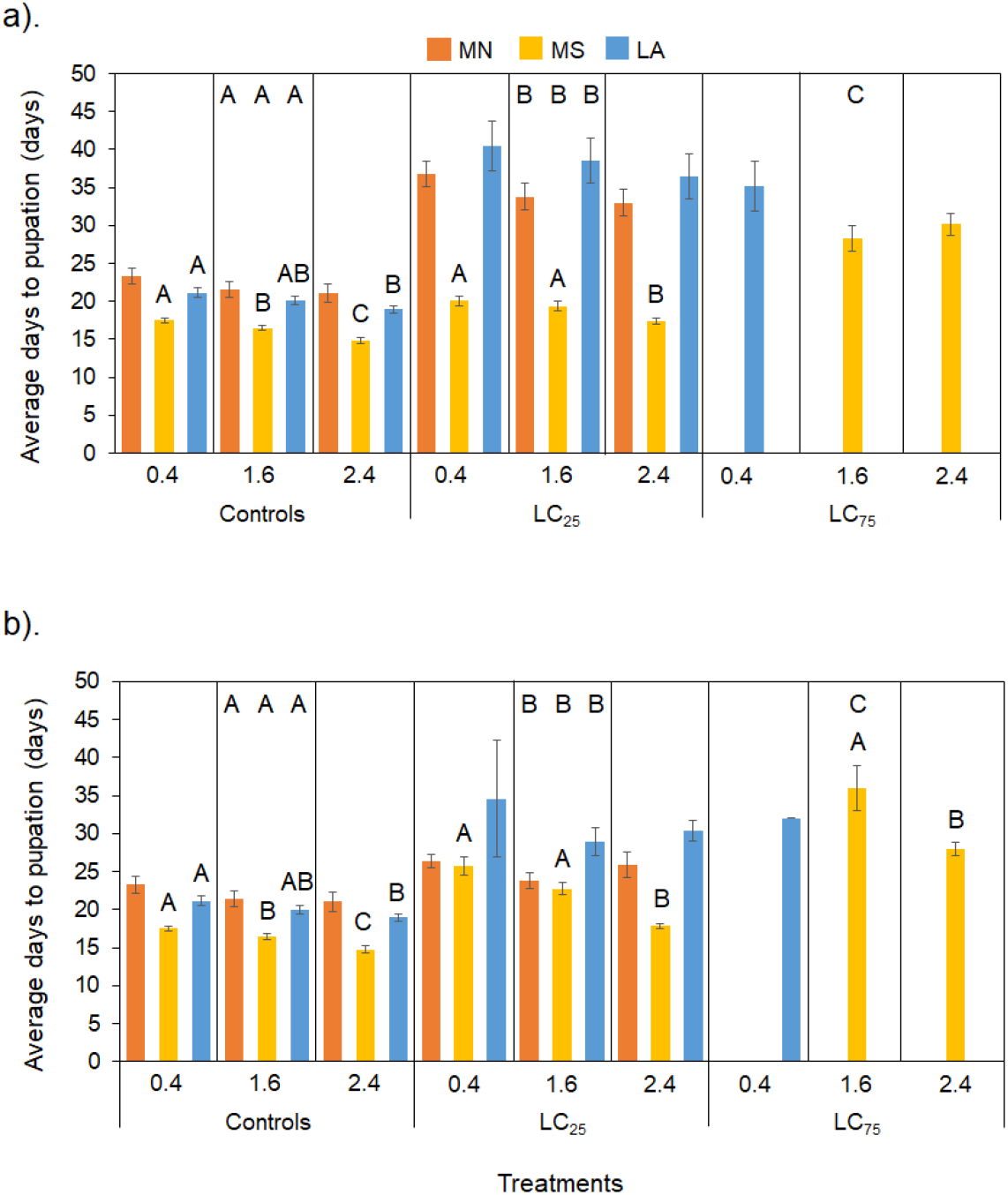
Average developmental time for all populations and Cry-diet treatments for Cry1Ac (a) and Cry1Ab (b). Error bars denote ± SE. Different letters at the top of graph indicate significant post-hoc differences between Cry treatments for each population, letters above bars indicate significant post-hoc differences between diets within each treatment at P ≤ 0.05.

**Figure 6.**
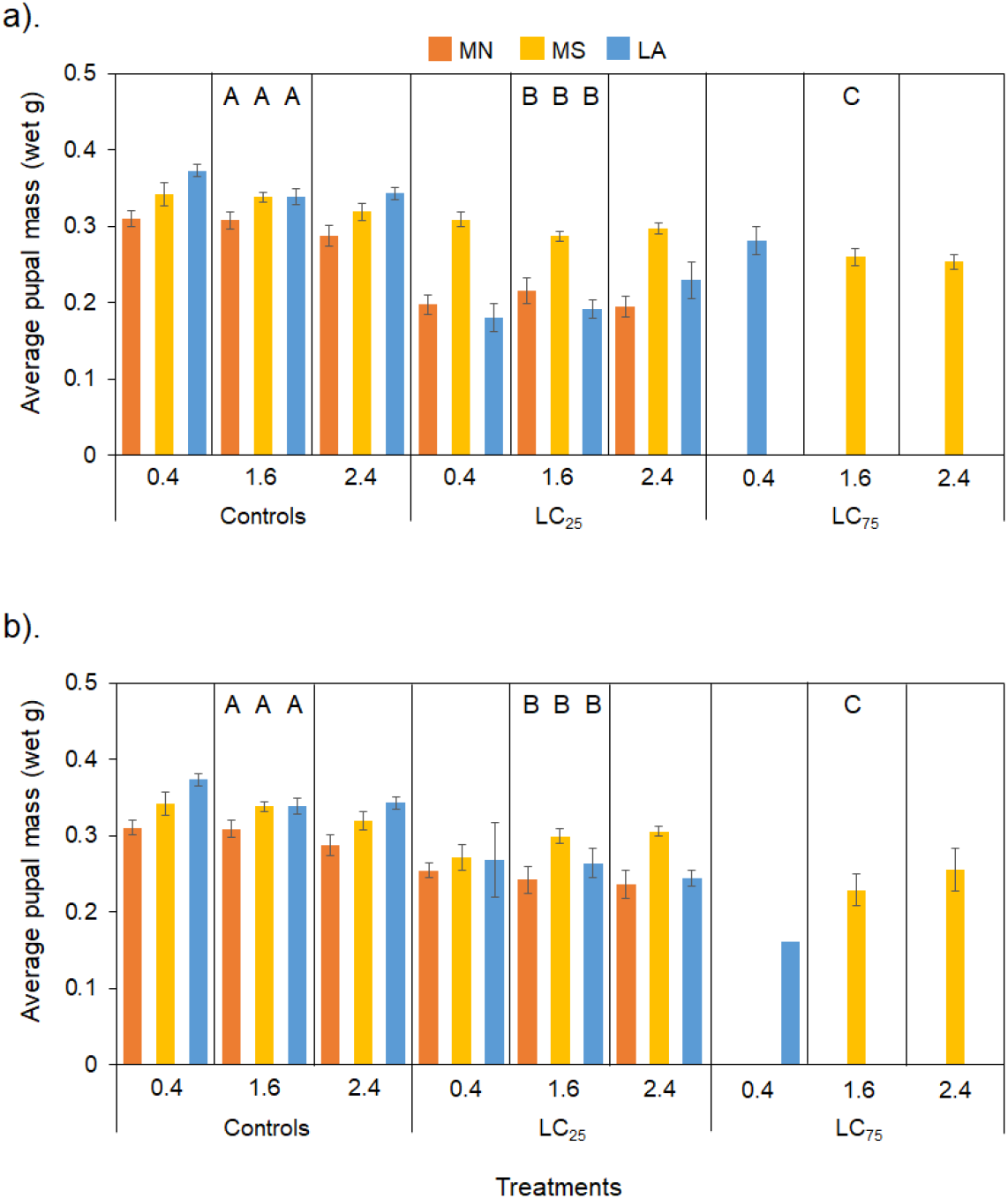
Average pupal mass for all populations and Cry-diet treatments. Error bars denote ± SE. Different letters at the top of graph indicate significant post-hoc differences between Cry treatments for each population at P ≤ 0.05.

### MS Population

Increasing concentrations of both Cry1Ac and Cry1Ab had significant negative effects on larval survival (Figure 3b,e,h; Figure 4b,e,h). For both Cry endotoxins, there was no statistical difference in survival between the controls and the LC_25_ treatments; however, survival was significantly lower in the LC_75_ treatment than both the controls and the LC_25_ treatment (Table 2). Average survival across diets in the control treatment was 89.6%. Based on the initial dose-response data for this population on the 1.6 diet, survival at the LC_25_ dose of Cry1Ac produced much higher than expected survival at 90.3%, which was only 3.5% lower than the survival for the 1.6 diet controls. Survival for the LC_75_ Cry1Ac treatment was also much higher than expected at 48.4%. For Cry1Ab, survival in both the LC_25_ and LC_75_ treatments was only slightly lower than expected, at 69.7% and 16.1% respectively.

The effect of diet on survival varied between Cry1Ac and Cry1Ab. For Cry1Ac, diet effects were only found in the control treatment (Table 3), with the protein-biased diet (2.4) producing significantly higher survival than the carbohydrate-biased diet (0.4) and the intake target diet (1.6) showing intermediate survival (Figure 3b). For Cry1Ab, significant diet effects were found in the controls and the LC_75_ treatment (Table 3). The control treatment was used for both Cry experiments; hence, the results were the same. All of the larvae fed the carbohydrate-biased diet (0.4) died at Cry1Ab LC_75_ concentration, while the 1.6 and 2.4 diets maintained significantly higher survival (Figure 4h).

Cry concentration had a significant impact on developmental time (Table 4), with similar trends for both Cry proteins. Figure 5 shows that days to pupation increased significantly as Cry exposure increased. The MS population was the only population to show pupation in the LC_75_ treatments and there were significant increases in developmental time between all concentrations for both Cry1Ac and Cry1Ab (Table 4). Diet also had a significant effect on developmental time. For Cry1Ac, diet effects were evident for the control and LC_25_ treatments. For controls, the 2.4 showed the shortest developmental time, followed by the 1.6, then the 0.4 diets. For the LC_25_ treatment, the protein-biased 2.4 diet again had the shortest time to pupation, while the 1.6 and 0.4 diets had similar but longer developmental times (Figure 5a). For Cry1Ab, control results were the same as for Cry1Ac because the same control group was used.

Diet effects were also seen for the LC_25_ and LC_75_ treatments (Table 5). At the LC_25_ exposure, the 0.4 and 1.6 were similar and produced longer developmental times than the protein-biased 2.4 diet.

No pupation was seen for the 0.4 diet at the LC_75_ exposure but developmental time was again significantly longer for the 1.6 diet treatment than the 2.4 diet treatment (Figure 5b). Cry concentration had a significantly effect on pupal mass for both Cry proteins (Table 6), with pupal mass decreasing significantly as Cry concentration increased (SI Table 2; Figure 6). Diet did not have a significant effect on pupal mass (Table 6).

#### LA Population

Increasing concentrations of both Cry1Ac and Cry1Ab had significant negative effects on larval survival (Figure 3c,f,i; Figure 4c,f,i). Table 2 shows that survival was significantly different across all Cry1Ac and Cry1Ab concentrations, with survival decreasing as concentrations increased. Average survival in the control treatment was high, at 91.7%. Based on the dose-response data for this population on the 1.6 diet, survival in the Cry1Ac LC_25_ and LC_75_ treatments was lower than expected at 47.8% and 0% respectively. Survival in the Cry1Ab LC_25_ treatment was remarkably accurate at 75.0%, while the LC_75_ treatment showed lower than expected survival at 0%.

There were minimal diet effects on Cry susceptibility for the LA population. Table 3 shows that significant diet effects were only evident for the Cry1Ac LC_75_ treatment. While mortality was 100% in all diet treatments at the LC_75_ concentration, larvae fed the protein-biased diet died more quickly than those fed either the 0.4 or 1.6 diet.

Cry concentration had significant effects on developmental time for both Cry proteins (Table 4), with time to pupation taking longer in the LC_25_ treatments than the controls (Figure 5). Diet had a significant effect on developmental time for the controls only (Table 5). Control treatments were the same for Cry1Ac and Cry1Ab experiments. Figure 5 shows that the protein-biased 2.4 diet produced the fastest developmental time, followed by the 1.6 diet, then carbohydrate-biased 0.4 diet, however, significant differences were only apparent between the 0.4 and 2.4 diets (SI Table 1). There was a significant effect of Cry concentration on pupal mass for both Cry proteins (Table 6). Figure 6 shows that pupal mass declined significantly as Cry concentration increased. No diet effect on pupal mass was found.

## Discussion

This study provides the first evidence for genetic-by-environment interactions between field-relevant macronutrient variability and Cry susceptibility in different populations of *H. zea*. By measuring larval susceptibility to Cry proteins using individuals from different geographical locations, we were not only able to assess the prevalence of nutritional impacts on the efficacy of *Bt* endotoxins, but also the role of gene-by-environment interactions in pest management. Deans et al. (2017) demonstrated that dietary macronutrients can have significant impacts on *H. zea*’s susceptibility to Cry1Ac in a laboratory strain; however, the current study shows that these impacts are strongly affected by genetic background and will likely vary in the field. Overall, our results indicate that interactions between diet and Cry susceptibility are incredibly complex and subject to the interplay between genetic and environmental factors.

The impact of Cry exposure on larval survival and performance was much more consistent across populations than the impact of diet on toxicity. The Cry bioassays showed that there were some initial differences in susceptibility between populations, but overall, lethal dose ranges overlapped considerably for all three populations. Although we couldn’t directly compare the Cry and diet results across populations due the fact that the experiments were done at different times, a qualitative comparison of the results show that the MS population had considerably higher survival than the other populations, as it was the only population with greater than 1 survivor in the LC_75_ treatments for both Cry proteins. The MS population also showed qualitatively higher performance in the face of Cry exposure. Although Cry had significant negative impacts on developmental time and pupal mass in all populations, the impacts in the LC_25_ treatments were noticeably smaller for the MS population (Figure 5 and 6). This is somewhat surprising given that the initial bioassays suggested that the LA population would likely have the highest tolerance. For Cry1Ac, the LA and MS populations showed greater than 25% and 75% mortality in the LC_25_ and LC_75_ treatments on the 1.6 diet, while the MS population had lower than expected mortality. For all populations, the results for Cry1Ab more closely matched the expected mortality values in the LC_25_ treatments, while morality was higher than expected for the LC_75_ treatments. Overall, this suggests that 10-day bioassays may underestimate mortality over the entirety of larval development.

The effect of diet on survival and performance differed between populations and, in some cases Cry protein, resulting in distinct patterns for each population. Diet had no impact on survival in the LC_25_ treatments, but a significant impact was apparent at the higher LC_75_ dose for all populations on at least one Cry protein. The MS population was unique in that a significant diet effect was evident in the control treatments as well. For the MN population, diet affected susceptibility to both Cry proteins, while it only impacted Cry1Ab susceptibility for the MS population and only Cry1Ac for the LA population. Additionally, the diets that showed optimal survival varied between populations. The intake target (1.6) and the protein-biased (2.4) diets showed similarly higher survival under Cry exposure for the MN and MS populations, while the intake target and carbohydrate-biased diet provided the highest survival for the LA population. Dietary effects on developmental time were also variable. There was no effect of diet on time to pupation for the MN population, an effect on both the controls and Cry treatments for the MS population, and only an effect on the controls for the LA population. Despite these differences, the protein-biased 2.4 diet treatments had the shortest developmental times in all cases.

Although we did not assess the genetic variability or relatedness between our populations, it is reasonable to assume that the differences we observed in Cry-diet interactions are largely the result of differences in genetic background, particularly since the environment was standardized for all populations. Even though *H. zea* is capable of migrating over long distances, strong differences were still observed between the LA and MS populations despite being separated by less than 300 km. In fact, the MN and LA populations had the most similar results, yet were the furthest apart. This is perhaps not surprising, however, as it is well-established that the ephemeral *H. zea* populations in MN are derived from seasonal migrations from populations in the Gulf (Sandstrom et al., 2007; Seymour et al., 2016). Genetic differences between populations likely reflect local adaptations to variations in exposure to Cry endotoxins, however, the MS and LA population were held in laboratory culture for a longer period than the MN population, so we are also unable to exclude genetic differentiation due to adaptation to laboratory conditions. In any case, the observed population-level differences do indicate that genetic background has a strong impact, not only on the relationship between nutrition and fitness under control conditions but also on the relationship between nutrition and Cry susceptibility. This suggests that nutritional state may impact Cry toxicity in multiple ways, perhaps acting at different points along the mode of action and likely implicating many different loci.

The results of Deans et al. (2017) showed that nutrition can mediate plasticity in *Bt* susceptibility by documenting variations in survival across different diet p:c ratios. The results of this study further indicate that there is likely a genetic basis for this plasticity, as variable nutritional effects were observed across different populations (Pigliucci, 2001). Despite this, the genetic mechanisms underlying these population-level differences remain unknown. Relevant differences between populations could be related to general differences in nutritional physiology or to differences in resource allocation to various processes used to mitigate toxicity. In Deans et al. (2017) deviations away from the self-selected *H. zea* intake target negatively affected Cry susceptibility; however, although the 1.6 diet was among those with the highest survival for all populations in this study, the MN and MS populations showed that the protein-biased also produced similarly high survival and the LA population showed that the carbohydrate-biased diet also produced similarly high survival. Furthermore, the 2.4 diet treatment, not the 1.6 diet, universally showed the best performance, with performance for the intake target diet being intermediate. It is possible that our populations may have local adaptations for intake targets that are different from that reported for the laboratory strain. Differences in nutrient regulation could explain some of the nutritional discrepancies observed across populations. If the actual intake target for the LA population were shifted to a more carbohydrate-biased p:c ratio, for instance, it may explain why survival was uniquely lower on the 2.4 diet in the LC_75_ Cry1Ac treatments for that population but not the others. Latitudinal changes in intake targets have been reported for field populations of grasshoppers, and it has been hypothesized that these changes may be tied to latitudinal temperature gradients (Parsons, 2011). Additional studies will be required to determine whether nutrient regulation varies across geographical populations and/or genetic backgrounds in *H. zea* and other insects.

Another possible explanation for these nutritional differences across populations may be related to differences in the physiological response to *Bt*. There are many points along the mode of action of *Bt* that can impact susceptibility. Several mechanisms of *Bt* resistance have been documented in a variety of lepidopteran species. Changes in gut pH can decrease solubility of the endotoxin, changes in the type and number of gut proteases can prevent formation of the protoxin, structural alterations in binding targets, such as cadherins, alkaline phosphatases, and membrane-bound aminopeptidases, can prevent pore formation, and increased stem cell activity can repair cellular damage, thereby reducing gut damage. All of these interventions have different physiological costs and, as such, may be constrained by nutrition in different ways. Given the high susceptibility of all our populations, genetic mutations affecting structural differences in binding sites are not likely relevant to this study, but more tractable interventions, like protease expression or stem cell activity, could vary across populations, eliciting different nutritional costs. For instance, some interventions may be more protein-intensive, while others energetically costly (carbohydrate-intensive). These physiological constraints could help to explain some of the differences between populations but additional work would be required to understand population-level responses to *Bt* and their constraints.

While the effects of genetic resistance are widely studied and consistently monitored in agroecosystems, the relative impact of gene-by-environment interactions on pest dynamics receive less attention. Genetic mutations that instantaneously alter the efficacy of insecticides are often easier to identify and characterize than failures due to plastic responses; however, gene-by-environment interactions could account for much of the inconsistencies observed in insecticide control. Because plant nutrient content is extremely variable through space and time, nutrition likely has important and widespread effects on herbivore pests. Plastic responses have often been disregarded because they are difficult to control under field conditions, but acknowledging their importance is necessary of effective pest control. For instance, Deans et al. (2016, 2017) demonstrated that failing to utilize field-relevant artificial diets, that is diets with macronutrient profiles that match what the insect is regulating for in the field, into insecticide resistance monitoring bioassays can significantly confound results.

Because our results suggest that different populations will be impacted by nutrition in different ways, we suggest that the macronutrient profiles of artificial diets used for baseline susceptibility bioassays and resistance monitoring should be customized to the pest and the cropping system. Ideally, bioassays should be run using an artificial diet matching the intake target, if one is available, as this most accurately reflects the nutritional state of insects in the field. However, polyphagous pests, such as *H. zea*, may not be able to reach their intake target in all crops. The macronutrient data for sweet corn (Deans et al., 2018) for instance, shows that all tissues contain a carbohydrate-biased p:c ratio. In this system, a more carbohydrate-biased artificial diet would be more appropriate to get an accurate view of susceptibility. Ultimately, more data on plant macronutrient content and insect nutritional ecology are needed to continue to optimize pest control strategies and overall resource management.

## Acknowledgements

This work was funded by a USDA BRAG grant #2015-33522-24099.

## Supplementary Tables/Figures

**SI Table 1.**
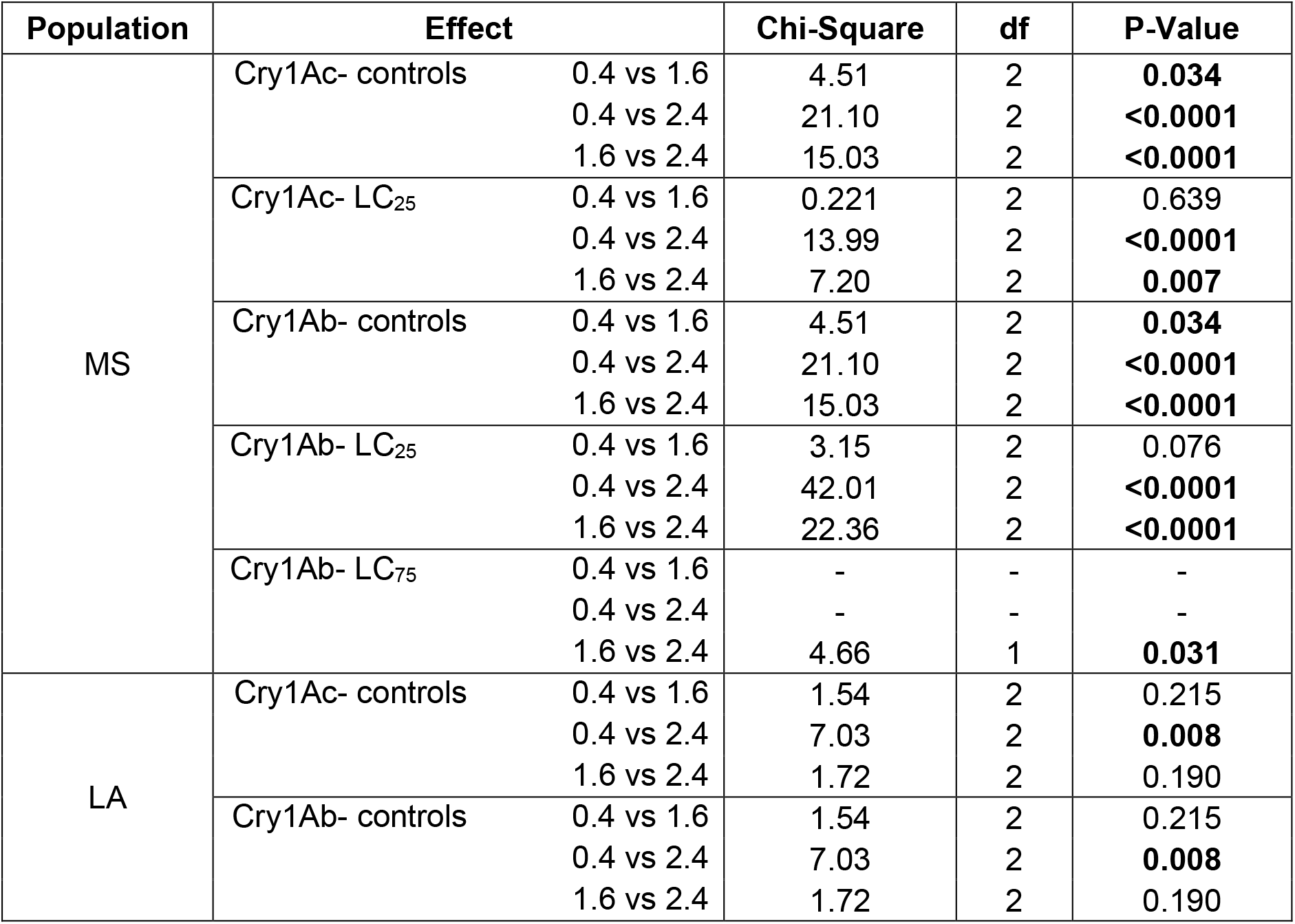
Post-hoc results for the effect of diet on developmental time. Bolded values indicate significance at P≤ 0.05

**SI Table 2.** Tukey HSD post-hoc results for the effect of Cry concentration on pupal mass for the MS population. There were less than 2 pupation events in the MN and LA populations so post-hoc analyses were not performed. Bolded values indicate significance at P ≤ 0.05.

| Population | Effect | P-Value |
| --- | --- | --- |
| MS | Cry1Ac 0 vs LC<sub>25</sub> | **<0.0001** |
|  | 0 vs LC<sub>75</sub> | **<0.0001** |
|  | LC<sub>25</sub> vs LC<sub>75</sub> | **<0.0001** |
|  | Cry1Ab 0 vs LC<sub>25</sub> | **<0.0001** |
|  | 0 vs LC<sub>75</sub> | **<0.0001** |
|  | LC<sub>25</sub> vs LC<sub>75</sub> | **0.035** |

## Notes

### Competing Interest Statement

The authors have declared no competing interest.

